# Photoperiod Sensitive Energy Sorghum Responses to Environmental and Nitrogen Variabilities

**DOI:** 10.1101/2020.11.03.366898

**Authors:** August Schetter, Cheng-Hsien Lin, Colleen Zumpf, Chunhwa Jang, Leo Hoffmann, William Rooney, D.K. Lee

**Affiliations:** Department of Crop Sciences, University of Illinois at Urbana-Champaign, Urbana, Illinois 61801, USA; Center for Advanced Bioenergy and Bioproducts Innovation, University of Illinois at Urbana-Champaign, Illinois 61801, USA; Carl R. Woese Institute for Genomic Biology, University of Illinois at Urbana-Champaign, Urbana, Illinois 61801, USA; Department of Soil and Crop Sciences, Texas A&M University, College Station, Texas, 77843, USA

**Keywords:** Photoperiod-sensitive sorghum, Environment, Feedstock quality, Nitrogen use efficiency

## Abstract

Recently introduced photoperiod-sensitive (PS) biomass sorghum (*Sorghum bicolor* L. Moench) needs to be investigated for their yield potentials under different cultivation environments with reasonable nitrogen (N) inputs. The objectives of this study were to 1) evaluate the biomass yield and feedstock quality of four sorghum hybrids with different levels of PS ranging from very PS (VPS) hybrids and to moderate PS (MPS) hybrids, and 2) determine the optimal N inputs (0~168 kg N ha^−1^) under four environments: combinations of both temperate (Urbana, IL) and subtropical (College Station, TX) regions during 2018 and 2019. Compared to TX, the PS sorghums in central IL showed higher yield potential and steady feedstock production with an extended daylength and with less precipitation variability, especially for the VPS hybrids. The mean dry matter (DM) yields of VPS hybrids were 20.5 Mg DM ha^−1^ and 17.7 Mg DM ha^−1^ in IL and TX, respectively. The highest N use efficiency occurred at a low N rate of 56 kg N ha^−1^ by improving approximately 33 kg DM ha^−1^ per 1.0 kg N ha^−1^ input. Approximately 70% of the PS sorghum biomass can be utilized for biofuel production, consisting of 58-65% of the cell wall components and 4-11% of the soluble sugar. This study demonstrated that the rainfed temperate area (e.g., IL) has a great potential for the sustainable cultivation of PS energy sorghum due their observed high yield potential, stable production, and low N requirements.

## Introduction

Herbaceous feedstock is a critical renewable resource for bioenergy production necessary to alleviate global dependence on fossil fuels [1]. Sorghum (*Sorghum bicolor* L. Moench), primarily known as a cereal or forage crop, is receiving increased attention as an energy crop for producing biofuel (e.g., ethanol). Based on its types and primary crop components, sorghum can be divided into several categories: grain sorghum with starch-rich grain, sweet sorghum with high soluble sugar, and forage sorghum with high leaf to stem ratios, silage sorghums and biomass sorghum (bioenergy) with high lignocellulosic components [2,3].

Biomass and sweet sorghum are considered as an ideal bioenergy crop for several reasons. Both types have excellent biomass yield potential, including energy-rich compositions such as structural and non-structural carbohydrates. In general, sorghum also has high tolerance to environmental stress (e.g., drought conditions, high salinity soils, or low soil fertility) and has the potential to sustain relatively high biomass production under adverse environments. Additionally, the long history of established sorghum production systems in the U.S. provides growers with the opportunity to quickly adopt a new crop into their existing production systems. Finally, the genetic improvement of sorghum through traditional and genomic approaches for multiple objectives has allowed for advantageous gains in yield potential, stress tolerance, and feedstock quality [3,4].

Plant breeders have utilized the knowledge of photoperiodism as a strategy to adjust sorghum maturity to maximimize the vegetative growth phase to improve biomass yield. These manifest as photoperiod-sensitive (PS) sorghum hybrids [4,5]. The PS sorghum hybrids remain in the vegetative growth phase until day length is less than a specific daylength; the daylength can vary depending on the genetics of the hybrid. As such, in temperate environments grain or seed production is essentially impossible because day lengths during the growing season (April – September) are much greater than the minimum required to initiate reproductive growth [4,6]. An extended period of vegetative growth provides sorghum with two significant advantages including 1) extending the duration of utilizing the solar radiation to cumulate biomass during the growing season, and 2) a high level of drought tolerance in rainfed cultivation environment [4,7,8]. For instance, the delayed maturity of the PS energy sorghum is usually associated with higher dry biomass yield ranging from 20 to 35 Mg ha^−1^ compared to grain sorghum (approximately two fold higher) [7,9,10]. Under good growing conditions (e.g., without water and nutrient deficiency), substantially increased biomass yields (e.g., >35 Mg DM ha^−1^) were also reported in TX [11–13].

In addition to high yield potential, biomass feedstock composition is also a crucial factor for biofuel conversion processes. Comprehensive understanding of the feedstock composition are important for multiple operational purposes, including storage, pre-treatments, and biorefinery operations [14,15]. Different sorghum types and hybrids contain specific compositional characteristics. For instance, the PS biomass sorghum hybrids tend to have concentrated high energy-dense structural compositions, such as glucans (26.9-31.8%), xylan (14.9-18.4%), and lignin (8.3-18.9%) [12,16]. Sweet sorghum hybrids usually have juicy stalks with high concentrations of non-structural carbohydrates [2,4]. McKinley et al. (2018) [12] reported that a new PS lignocellulosic energy sorghum hybrid can contain up to 60 % of fermentable components (dry biomass), including 50 % of structural cellulose, xylan, galactan, arabinan, and 10% of non-structural water-soluble sugars, that can be utilized for ethanol production via biochemical conversion processes. Carbon-rich lignin, although undesirable for biochemical conversion, can be accounted for another 10% and utilized for producing biofuel such as crude bio-oil using the thermochemical conversion techniques, such as fast pyrolysis. In contrast, ash (inorganic compounds) content is a hindrance to all conversion processes not only because ash has limited energy potential but likely reduces the conversion effectiveness resulting from strong catalytic effects on the processing pipelines [15,17,18].

Variations in both biomass yield and compositions of sorghum hybrids are also substantially influenced by growing conditions and management practices [12,16,19–23]. Gill et al. (2014) [21] reported that the production environment affected sorghum biomass yield; yields ranged from 1.5 to 41.1 Mg DM ha^−1^ across seven locations in the U.S. (i.e., Kansas, Iowa, Kentucky, Mississippi, North Carolina, and College Station and Corpus Christi in Texas) over a five years (2008-2012). The aboveground dry biomass likely increased by improving management practices, such as adequate nitrogen (N) input and irrigation systems [7,10,20,24]. Also, environmental variability substantially influenced biomass compositions. Packer (2011) [25] examined the compositional variability in 15 PS sorghum hybrids in five environments based on different soil types and climate regions across TX and showed that the concentrations of cellulose ranged from 26.9 to 31.8%, xylan from 14.9 to 18.4%, and lignin from 8.3 to 18.9%. For the field management effects, McKinley et al. (2018) [12] reported that the irrigated plots increased the concentration of the overall cell-wall composition of the PS hybrids by 20% compared to the non-irrigated fields. Although Amaducci et al. (2004) [20] did not observe a significant management effect, including nutrient input and tillage practice, on biomass compositions, Almodares et al. (2009) [26] showed that the increased N input reduced both soluble carbohydrates and fiber contents.

Many studies indicated that N application is the predominant energy input for the lignocellulosic energy crops among other management operations [27–29]. Compared to grain sorghums and other biofuel feedstock resources (e.g., sugarcane and maize), biomass sorghum requires less N input to accumulate lignocellulosic components. The lower N requirement implies that biomass sorghum can minimize the production cost and the environmental impact while maximizing biomass yield [26,30]. Olsen et al. (2013) [31] showed that the PS biomass energy sorghum had higher N use efficiency (NUE) than other candidate bioenergy species (e.g., grain and sweet sorghums, corn (*Zea mays*), and switchgrass (*Panicum virgatum*)) and the NUE likely improved by increasing vegetative growth duration. Nitrogen response of biomass sorghum, however, varies with the growing conditions, cultivar type, and cropping system, including the expected yield, phenotypic characteristics (e.g., plant height, stem thickness, and leaf to stem ratio), and biomass compositions [7,10,31]. For new energy sorghum hybrids, it is critical to investigate the effect of N practices on sorghum yield potential and feedstock composition across environmental gradients for management optimization. Therefore, the overall goal of this study was to determine the optimum management practice for sustainable biomass feedstock production of bioenergy sorghum. The specific objectives were to 1) evaluate the yield potential of four PS sorghum hybrids, and 2) determine the optimum N rate by evaluating the effect of N fertilization on energy sorghum biomass yield, feedstock compositional characteristics, and N removal and use efficiency in two growing environments, Illinois and Texas.

## Materials and Methods

### Research Sites and Experimental Design

A field study was conducted at the Energy farm of the University of Illinois located in Urbana, Illinois (IL, 40°3'N, 88°12'W) and the Texas AgriLife Research Farm near College Station (ARECCS), TX (30°32'N, 94°26'W) during the growing seasons of 2018 and 2019 (Table 1). A temperate humid continental climate type was categorized for the IL location with an average annual temperature of 11.5 °C and annual precipitation of 1,030 mm for the past 30 years (1990-2019 shown in Table 1). The soil type of the main plot in IL was predominated by Drummer silty clay loam (fine-silty, mixed, superactive, mesic Typic Endoaquolls). The TX location was categorized as the subtropical climate type with an average annual temperature of 20.7 °C and annual precipitation of 1,020 mm (30-year average). The primary soil type in TX was dominated by Ships clay loam (very-fine, mixed, active, thermic Chromic Hapluderts). Table 2 showed additional soil-related information in both locations and years. The IL site typically has longer daylengths than the TX site during the growing season (Fig. 1). The local monthly precipitation and temperature in 2018 and 2019, along with thirty-year averages (1990-2019) were obtained from National Oceanic and Atmospheric Administration for the IL site (Champaign-Urbana Willard Airport, USW00094870) and the TX site (College Station Easterwood Field, USW00003904).

**Table 1.** The information of the experimental locations, environmental conditions, and field activities, including dates for planting, and N fertilizer application.

| Location | 30-year average (annual) |  | Year | Planting | N appl | Harvest |  | Growth period (days) |  | Cumulative daylength (hrs) |  | GDD |  |
| --- | --- | --- | --- | --- | --- | --- | --- | --- | --- | --- | --- | --- | --- |
|  | Precip (mm) | Temp (°C) |  |  |  | VPS | MPS | VPS | MPS | VPS | MPS | VPS | MPS |
| Urbana, IL<br>(40° 3'N, 88°12'W) | 1030 | 11.5 | 2018 | May 18 | May 22 | Oct. 09 | Sep. 13 | 144 | 118 | 1998 | 1687 | 1754 | 1470 |
|  |  |  | 2019 | Jun. 01 | Jun. 07 | Oct. 09 | Sep. 19 | 130 | 110 | 1793 | 1556 | 1509 | 1314 |
| College Station, TX<br>(30° 32'N, 94°26'W) | 1020 | 20.7 | 2018 | May 03 | May 03 | Sep. 20 | Sep. 20 | 140 | 140 | 1898 | 1898 | 2451 | 2451 |
|  |  |  | 2019 | May 16 | May 16 | Sep. 17 | Sep. 17 | 124 | 124 | 1685 | 1685 | 2219 | 2219 |

Precip: Annual precipitation
Temp: Annual mean temperature
N appl: N fertilizer application
VPS: Very photoperiod sensitive
MPS: Moderate photoperiod sensitive
GDD: Growing degree days with a base temperature of 11°C accumulated during the growth period

**Table 2.** Soil characteristics at the depths of 0-15 and 15-30 cm for each growing environment, including two experimental sites of IL and TX during the growing seasons of 2018 and 2019.

| Location<br>(soil types) | Year | Depth<br>(cm) | Soil pH | Organic Matter<br>(%) | CEC<br>(meq 100g<sup>-1</sup>) | P<br>(kg ha<sup>-1</sup>) | K<br>(kg ha<sup>-1</sup>) |
| --- | --- | --- | --- | --- | --- | --- | --- |
| Urbana, IL<br>(Drummer silty clay loam) | 2018 | 0-15 | 6.4 | 5.4 | 20.0 | 80.7 | 197.3 |
|  |  | 15-30 | 6.4 | 4.7 | 20.7 | 35.9 | 141.2 |
|  | 2019 | 0-15 | 6.5 | 4.9 | 19.6 | 100.9 | 246.6 |
|  |  | 15-30 | 6.9 | 4.1 | 19.1 | 58.3 | 179.3 |
| College Station, TX<br>(Ships clay loam) | 2018 | 0-15 | 8.0 | 3.4 | 54.4 | 103.1 | 892.2 |
|  |  | 15-30 | 8.0 | 3.5 | 56.2 | 31.4 | 625.4 |
|  | 2019 | 0-15 | 8.0 | 3.4 | 53.0 | 94.2 | 887.7 |
|  |  | 15-30 | 8.1 | 3.0 | 53.4 | 49.3 | 668.0 |
| Methods | - | - | Water | Loss on ignition | - | Mehlich 3 | Mehlich 3 |

**Figure 1.**
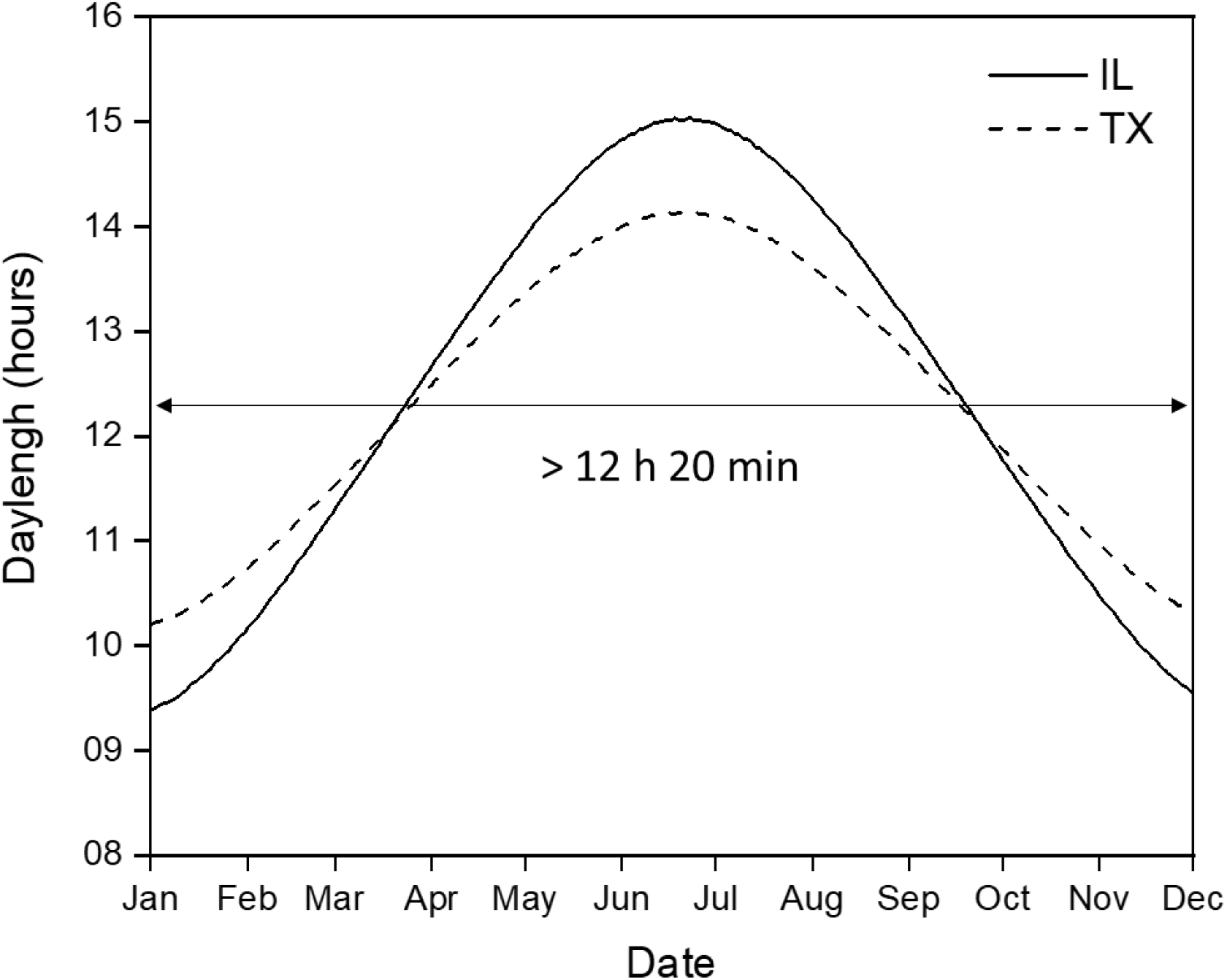
Daylight hours at Urbana (IL) and College Station (TX) in 2019.

Four hybrids of the PS energy sorghum were selected for this study: TX08001, TX17500, TX17600, and TX17800. The TX08001 and TX17500 hybrids initiate reproductive growth when day lengths are less than 12’20” and are categorized as very PS (VPS) hybrids; these hybrids will not flower in IL because freezing temperatures occur before flowering would occur. Compared to VPS hybrids, the TX17600 and TX17800 are less sensitive to this daylength threshold and categorized as moderate PS (MPS) hybrids with reproductive growth is initiated when daylengths pass below 12’40” to 13’00”. The phenotypic characteristics of four hybrids collected from College Station, TX are also shown in Table 3. Generally, four PS hybrids have a good standability with tall stalks ranging from 3 to 5 m. The four hybrids have different stalk thickness at the order of TX08001 and TX17800 (thick) > TX17600 (medium) > TX17500 (thin). Both TX17500 and TX17600 have juicy stalks, and TX17800 has a dry stalk. Four N treatments with the application rate of 0, 56, 112, and 168 kg-N ha^−1^ were evaluated for the biomass response. The experimental design was the split-plot using N treatments as the main plot and hybrids as the sub-plot design within a randomized complete block with four replications. Each subplot was 3 m by 12 m, consisting of 8 rows with 0.76 m row spacing.

**Table 3.** Phenotypic information of four photoperiod-sensitive energy sorghum hybrids.

| Maturity | Hybrid | Standability | Height (m) | Stalk thickness (mm) | Stalk Composition |
| --- | --- | --- | --- | --- | --- |
| VPS | TX08001 | Good | Tall | Thick | Intermediate |
|  | TX17500 | Excellent | Tall | Thin | Juicy |
| MPS | TX17600 | Good | Tall | Medium | Juicy |
|  | TX17800 | Good | Tall | Thick | Dry |

VPS: Very photoperiod sensitive
MPS: Moderate photoperiod sensitive

The summary of field operations was described in Table 1. In IL, the soybean stubble was disked prior to planting, and sorghum was planted at 180,000 seeds ha^−1^ using an ALMACO four-row Kinze planter (Nevada, IA). In TX, the prior crop was cotton; fields are cultivated and prepared in the fall and sorghum was planted at 180,000 seeds ha^−1^ using an ALMACO four-row Max-emerge planter. Both locations had the same N fertility program, and aqueous urea ammonium nitrate (UAN: 32-0-0) was knife-injected approximately a week after planting in IL and at the same day after planting in TX. The biomass harvest timing was based on site observation, when 50% of sorghum plants displayed visible inflorescences (heading). For both locations, the plots were harvested by cutting 4 rows in the middle of each plot using a John Deere (Moline, IL) forage harvester with a TCI Research Plot Sampler 130S (Williamstown, WI).

### Plant Tissue and Chemical Composition Analysis

Biomass tissue samples were directly collected from the TCI Research Plot Sampler 130S (Williamstown, WI) during harvest, and fresh sample weight was recorded. The biomass subsamples were dried in an oven for three days at 60°C to measure dry matter content and saved for tissue chemical composition analysis. Dried tissue samples were ground through a 2 mm screen in order to homogenize the sample and two subsamples were collected. One sample was analyzed for total N by a dry combustion method using a LECO FP-528 N/Protein Determinator (Leco Inc., St. Joseph, MI), while phosphorus (P), and potassium (K) were analyzed using a Perkin-Elmer Optima 8300 inductively coupled plasma (ICP) spectroscopy (PerkinElmer, Inc., Waltham, MA) following a concentrated HNO_3_ and HCl microwave digestion procedure using a MARSXpress vessel (CEM, Matthews, NC). Nutrient and mineral removal by sorghum biomass was calculated by multiplying biomass yield by the corresponding concentration of each element (kg ha^−1^). The other samples were analyzed for feedstock chemical composition using the near-infrared spectroscopy (NIRS) compositional prediction model developed by the National Renewable Energy Laboratory, and the analytical process was modified based on the published protocol [32]. The NIRS analysis provides a more detailed insight on biomass composition, including structural glucan, xylan, galactan, arabinan, acetyl, protein, and non-structural ash, water (e.g. sucrose) and ethanol extractives (e.g., chlorophyll and waxes).

### Nitrogen Use Efficiency

The entire plant aboveground biomass was harvested for the estimations of biomass yield, N uptake/removal, and N use efficiency. Physiological N use efficiency (PNUE) was determined by a ratio of DM biomass yield (BY) to the total N uptake (TNU), shown on Eq. 1 [31,33,34]. The increased biomass yield per applied unit of N fertilizer was defined as the N input efficiency (NIE), which is often referred to the agronomic NUE and was calculated based on Eq. 2. The amount of N in crop biomass (or crop nitrogen removal) that was originated from the applied N was defined as the N recovery efficiency (NRE) shown in Eq. 3.

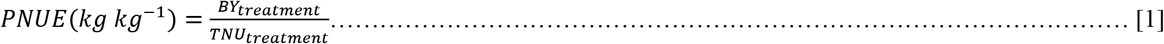

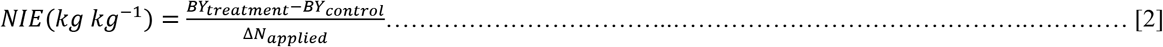

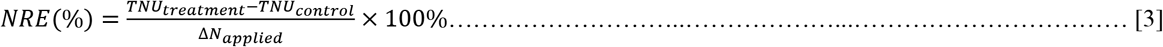

BY_treatment_ and BY_control_ (or TNU_treatment_ and TNU_contrtol_) are BY (or TNU) for different N treatments and control plots (zero N input), respectively. The ΔN_applied_ is the difference in UAN-N applied between treatments and control [35].

### Statistical Analysis

Statistical analysis with PROC MIXED procedure in SAS [36] was used to test the effect of locations (IL and TX), harvest year (2018 and 2019), N rate (four rates), hybrids (four hybrids) and their interactions on biomass yield production, plant tissue nutrient concentrations and removal, NUE, and feedstock chemical compositions using ANOVA. Locations, hybrid, N rates, harvest year, and their interactions were considered as fixed effects, where replications and their interactions were considered random effects. All significant difference was determined at P ≤ 0.05. Pairwise mean comparisons were made using the Fisher’s least significant difference (LSD) test.

## Results

### Environment

Monthly precipitation and temperature during the study period (2018 and 2019) and their 30-year average (1990-2019) for two experimental sites (i.e., Urbana, IL, and College Station, TX) are shown in Figure 2. During the growing season from May (planting) to Oct (harvest), the cumulative precipitation was higher in 2018 than 2019 and the 30-year average in both locations (IL: 676-mm in 2018 and 504-mm in 2019; TX: 657-mm in 2018 and 527-mm in 2019). The monthly temperature was similar between 2018 and 2019 for both IL and TX. Compared to the 30-year averages, the precipitation pattern showed more fluctuations than the temperature. In TX, the cumulative precipitation during the growing season (May-Oct) was predominated by the precipitation in Sept (210-mm) and Oct (298-mm) in 2018, accounting for 77% (508-mm out of 657-mm); by contrast, the Sept and Oct precipitation only contributed approximately 27% to the May-Oct cumulative precipitation (142-mm out of 527-mm) in 2019. This variation is typical in Texas and is associated with tropical storms from August to October. In IL, the sum of precipitation in Sept and Oct accounted for 24% and 28% of the overall precipitation during the growing season in 2018 and 2019, respectively, both of which showed the similar trend to the 30-average results (i.e. a total of 164-mm in Sept and Oct out of 600-mm from May to Oct). The soil chemical properties were also different in both locations. The TX site showed higher soil pH (8.0-8.1), CEC (53.0-56.2 meq 100g^−1^), and K content (625.4-892.2 kg ha^−1^) but lower organic matter (3.0-3.5%) than the IL site having soil pH ranging from 6.4 to 6.9, CEC from 19.1 to 20.7 meq 100g^−1^, soil K from 141.2 to 246.6 kg ha^−1^, and organic matter from 4.1 to 4.9% (Table 2).

**Figure 2.**
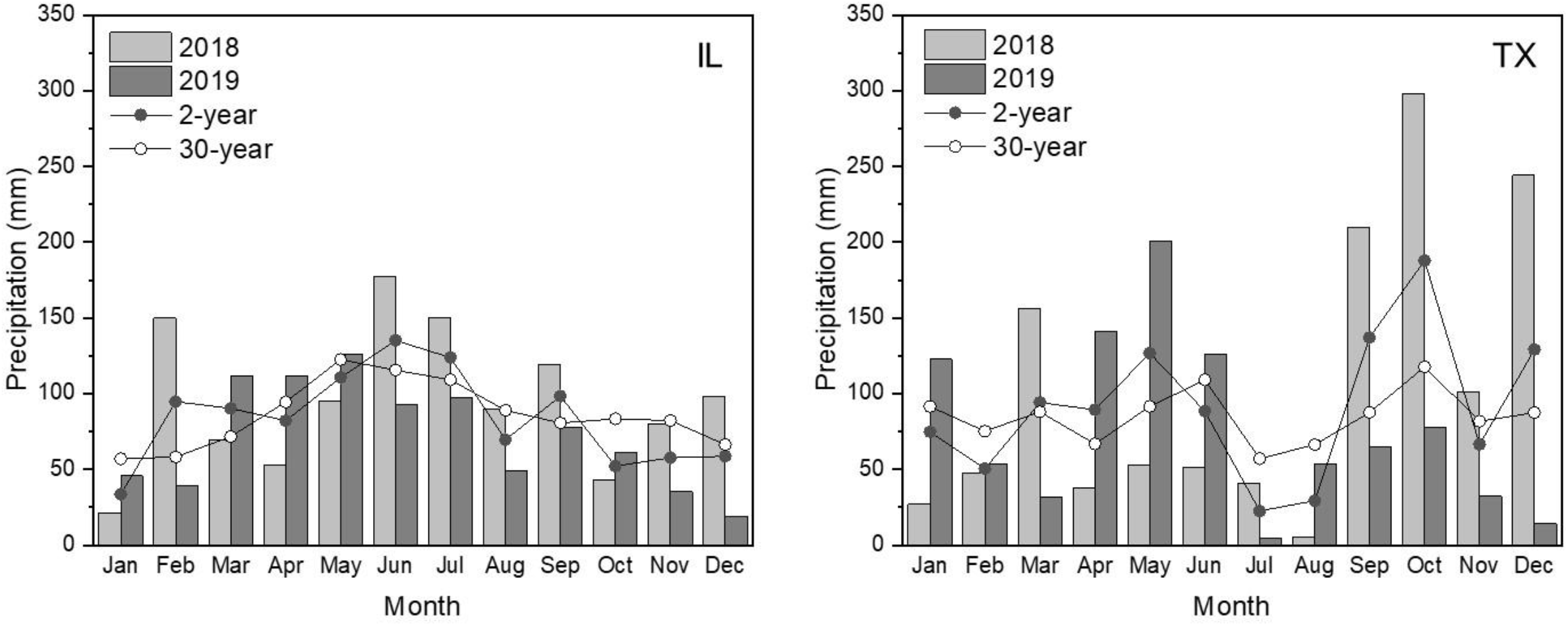
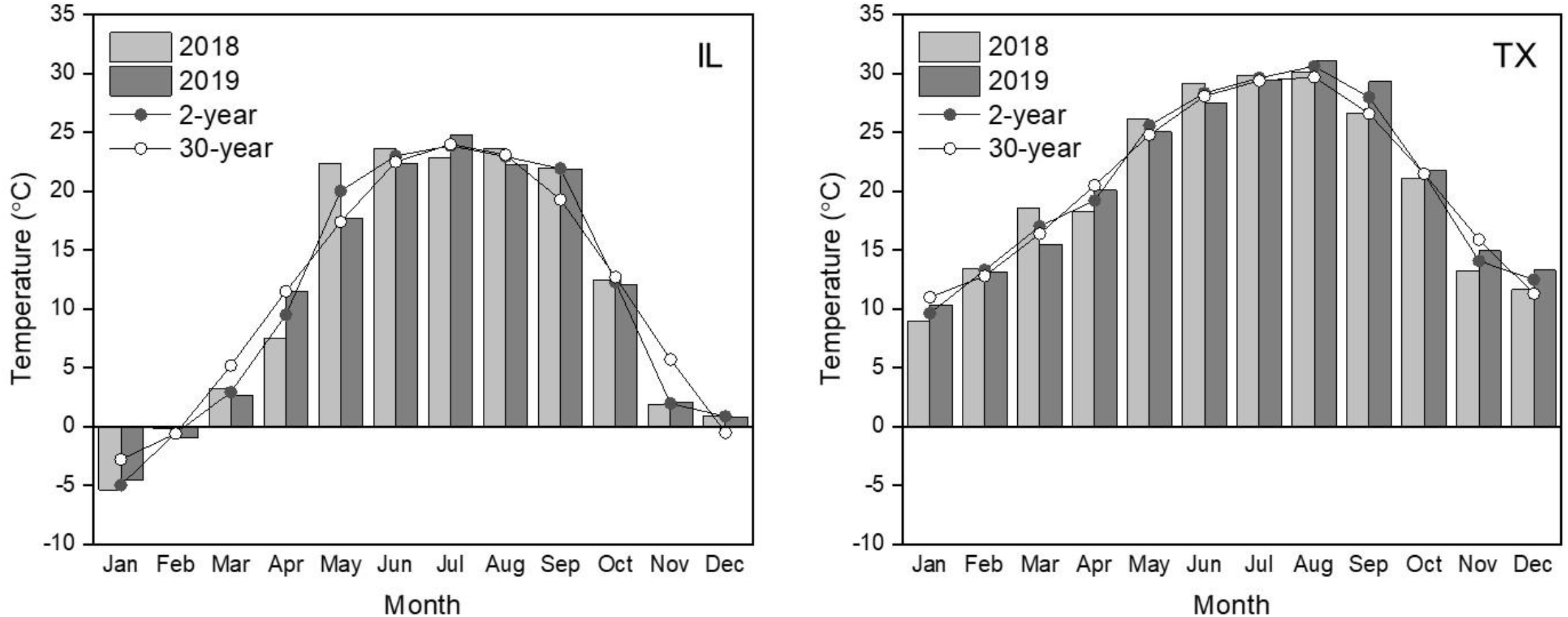
Local weather conditions in the experimental site across the two years of study including (a) monthly average precipitation and (b) monthly temperature and the 30‐year monthly average (1990-2019) (data: NOAA).

### Biomass Yield

The effects of four factors (location, year, N-rate, and hybrid) on biomass yield are shown in the analysis of variance (ANOVA) results (Table 4). The four-way interaction among these factors was not significant for biomass yield; however, the three-way interaction of location, year, and hybrid significantly influenced sorghum biomass yield (Table 4). The three-way interaction was caused by low biomass yield and variation among hybrids in TX-2018 (Fig. 3a). Except for TX17600, the yields of TX08001, TX17500, and TX17800 were significantly lower in TX-2018 compared to the other three environments (IL-2018, IL-2019, and TX-2019). Contrasted to yield averaged across IL-2018, IL-2019, and TX-2019, the biomass yield was lower by 21%, 47%, and 21% for TX08001, TX17500, and TX17800, respectively, in TX-2018. The two-way interaction between location and hybrid also affected biomass yield (Table 4), and the 2-year average showed that TX08001, TX17500, and TX17800 hybrids had higher yield potential in IL than TX except for TX176000 (Table 5). In IL, the TX08001, TX17500, and TX17800 biomass yield averaged across 2018 and 2019 were significantly higher by 10%, 22%, and 15 %, respectively, compared to the yield from TX. The average across the two years and four hybrids showed that the PS sorghums had higher yield potential in IL (20.4 Mg DM ha^−1^) than TX (19.0 Mg DM ha^−1^) (Table 5). The interaction between N rate and other factors did not show any significant impact on biomass yield (Table 4). Increased N rate generally increased the sorghum biomass yield (Table 4). However, the biomass yields of four hybrids showed similar responses to the increased N rate, and there was no benefit of N rates above 112 kg ha^−1^ (Table 6).

**Table 4.** Analysis of variance (ANOVA) showed the effects of main factors of location (L), year (Y), N-rate (Nr), and hybrids (H) and their interactions on biomass yield, biomass nutrient concentration, nutrient removal, and indices of physiological N use efficiency (PNUE), N input efficiency (NIE), and N recovery efficiency (NRE) with a significance level of 0.05.

| Effect | Yield | Plant nutrient concentration |  |  | Plant nutrient removal |  |  | Efficiency index |  |  |
| --- | --- | --- | --- | --- | --- | --- | --- | --- | --- | --- |
|  |  | N | P | K | N | P | K | PNUE | NIE | NRE |
| L | 0.0018* | 0.1466 | <.0001* | <.0001* | 0.0591 | <.0001* | <.0001* | 0.1081 | 0.5924 | 0.6553 |
| Y | <.0001* | 0.0461* | <.0001* | 0.7168 | <.0001* | 0.3914 | <.0001* | 0.0157* | 0.0015* | 0.0050* |
| Nr | 0.0009* | 0.0098* | 0.1846 | 0.9016 | 0.0015* | 0.0807 | 0.0936 | 0.0133* | 0.1111 | 0.7951 |
| H | 0.0003* | 0.0064* | <.0001* | 0.0016* | 0.1081 | 0.4298 | 0.1117 | 0.0098* | 0.1019 | 0.6568 |
| L x Y | <.0001* | <.0001* | 0.0090* | <.0001* | 0.1246 | <.0001* | <.0001* | <.0001* | 0.0020* | 0.5527 |
| L x Nr | 0.7042 | 0.0148* | 0.2967 | 0.0061* | 0.0089* | 0.0839 | 0.0394* | 0.0759 | 0.9336 | 0.1369 |
| L x H | <.0001* | 0.0036* | 0.1309 | 0.0406* | 0.1987 | 0.0123* | <.0001* | 0.0096* | 0.0604 | 0.1642 |
| Y x Nr | 0.1421 | 0.9391 | 0.1085 | 0.2448 | 0.1934 | 0.0821 | 0.0009* | 0.9708 | 0.4437 | 0.6996 |
| Y x H | 0.0001* | 0.020* | 0.0040* | 0.0006* | 0.0142* | 0.5053 | 0.0159* | 0.0350* | 0.4846 | 0.3117 |
| H x Nr | 0.2865 | 0.9829 | 0.5137 | 0.0253* | 0.9361 | 0.895 | 0.1392 | 0.9010 | 0.9452 | 0.9809 |
| L x Y x Nr | 0.1823 | 0.0940* | 0.581 | 0.1947 | 0.8557 | 0.8916 | 0.073 | 0.1717 | 0.4873 | 0.8456 |
| L x Y x H | 0.0020* | 0.0016* | 0.0362* | 0.0009* | 0.2818 | 0.1659 | 0.0140* | 0.0145* | 0.0752 | 0.4543 |
| L x Nr x H | 0.6587 | 0.9374 | 0.7412 | 0.2127 | 0.5951 | 0.6556 | 0.5299 | 0.9295 | 0.9637 | 0.8999 |
| Y x Nr x H | 0.8655 | 0.8764 | 0.2792 | 0.0635 | 0.6477 | 0.8527 | 0.0676 | 0.7576 | 0.8567 | 0.9710 |
| L x Y x Nr x H | 0.3477 | 0.9025 | 0.0893 | 0.291 | 0.8725 | 0.4621 | 0.0986 | 0.9223 | 0.3761 | 0.9115 |

\* Significant effect at the level of $P < 0.05$

**Table 5.** The significant effect of location (IL and TX) x hybrid (TX08001, TX17500, TX17600, and TX17800) on dry biomass yields, nutrient concentrations, and removal (N, P, and K). Lowercase letters indicate mean separation α=0.05 organized highest to lowest value for each column (no mean separations were applied if the variable effect was not significant).

| Location | Hybrid | Yield<br>(Mg ha<sup>-1</sup>) | Nutrient concentration (g kg<sup>-1</sup>) |  |  | Nutrient removal (kg ha<sup>-1</sup>) |  |  |
| --- | --- | --- | --- | --- | --- | --- | --- | --- |
|  |  |  | N | P | K | N | P | K |
| IL | TX08001 | 20.9ab | 7.3d | 1.4 | 11.7d | 149.9 | 29.4ab | 243.7de |
|  | TX17500 | 20.2bc | 7.7bcd | 1.6 | 13.2c | 158.4 | 31.6a | 261.9d |
|  | TX17600 | 19.5bcd | 8.4abc | 1.4 | 10.5de | 159.9 | 27.8bc | 203.5f |
|  | TX17800 | 20.9ab | 8.3abc | 1.3 | 10.2e | 175.6 | 27.7bc | 213.5ef |
| TX | TX08001 | 18.9cd | 7.7bcd | 1.3 | 20.4ab | 142.7 | 24.2cde | 384.4b |
|  | TX17500 | 16.5e | 9.2a | 1.4 | 20.3ab | 136.3 | 21.9e | 323.9c |
|  | TX17600 | 22.5a | 7.6cd | 1.2 | 18.9b | 167.0 | 26.3bcd | 427.8a |
|  | TX17800 | 18.3d | 8.5ab | 1.3 | 20.6a | 147.3 | 23.0de | 367.6b |
| Location<br>mean | IL | 20.4a | 7.9 | 1.4a | 11.4b | 160.9 | 29.1a | 230.6b |
|  | TX | 19.0b | 8.3 | 1.3b | 20.1a | 148.3 | 23.9b | 375.8a |

**Table 6.** The significant 2-way interaction between location (IL and TX) and N-rate (0, 56, 112, and 168 kg-N ha^−1^) on biomass yields, nutrient concentration, and removal. Lowercase letters indicate mean separation α=0.05 organized highest to lowest value (no mean separations were applied if the location x N rate interaction was not significant).

|  | IL |  |  |  | TX |  |  |  |
| --- | --- | --- | --- | --- | --- | --- | --- | --- |
| N rate | 0 | 56 | 112 | 168 | 0 | 56 | 112 | 168 |
| ----- Dry biomass (Mg ha<sup>-1</sup>) ----- |  |  |  |  |  |  |  |  |
| Yield<sup>†</sup> | 18.1 | 19.8 | 21.5 | 22.1 | 17.1 | 18.9 | 20.1 | 20 |
| ----- Biomass nutrient concentration (g kg<sup>-1</sup>) ----- |  |  |  |  |  |  |  |  |
| N | 6.9d | 7.2d | 7.8bcd | 9.5a | 7.5cd | 8.5abc | 8.3bc | 8.7ab |
| P<sup>†</sup> | 1.4 | 1.5 | 1.4 | 1.4 | 1.4 | 1.4 | 1.2 | 1.2 |
| K | 12.1bc | 12.3b | 10.7bc | 10.6c | 19.5a | 19.5a | 20.2a | 21.0a |
| ----- Plant nutrient removal (kg ha<sup>-1</sup>) ----- |  |  |  |  |  |  |  |  |
| N | 126.7d | 141.7cd | 166.2b | 209.2a | 125.1d | 149.2bcd | 159.6bc | 159.3bc |
| P<sup>†</sup> | 25.8 | 29.4 | 29.7 | 31.6 | 23.2 | 24.8 | 24.8 | 22.6 |
| K | 217.9c | 242.5c | 227.6c | 234.4c | 330.4b | 365.5ab | 402.9a | 404.3a |

†: no significant location x N rate interaction effect

**Figure 3.**
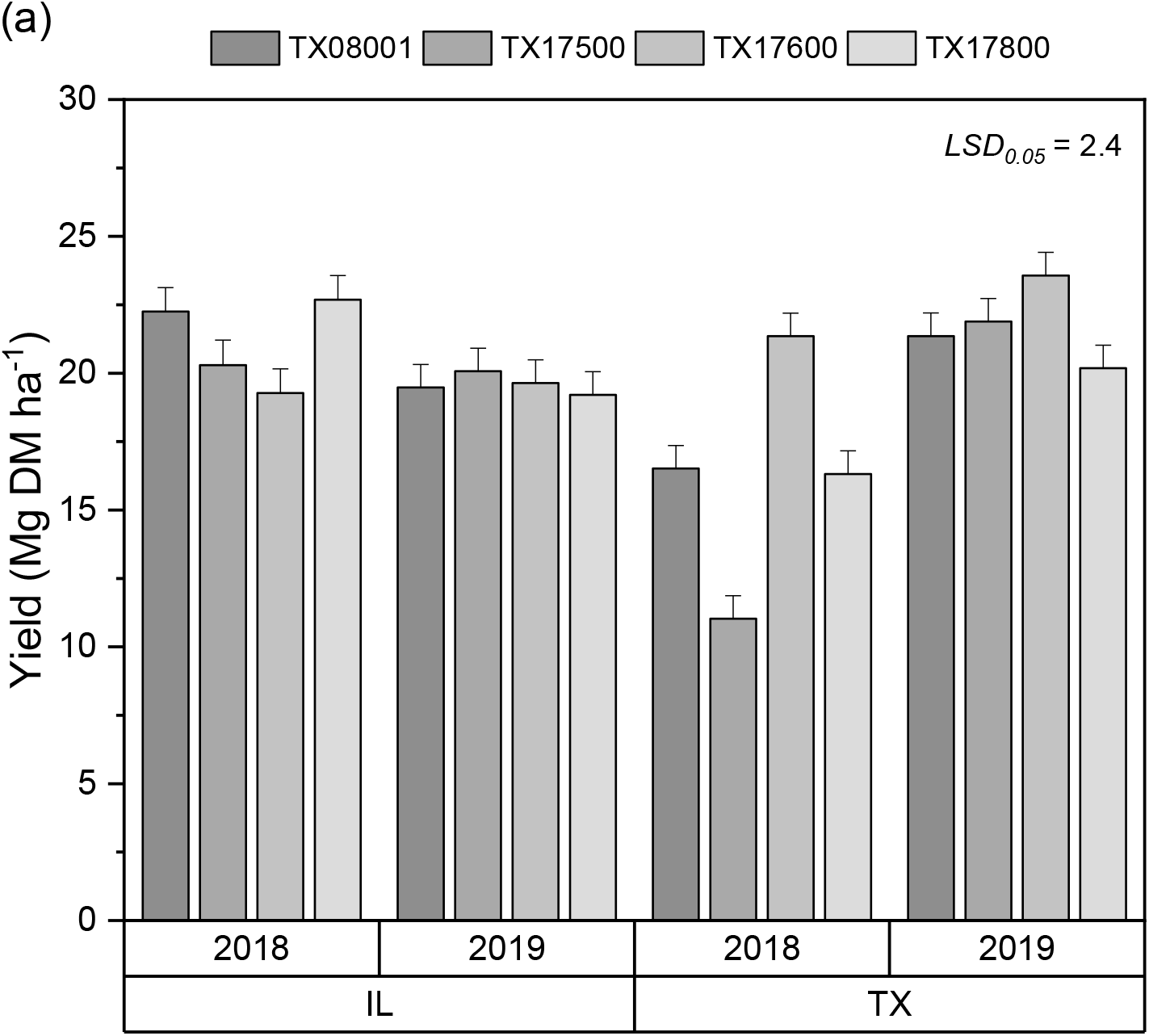
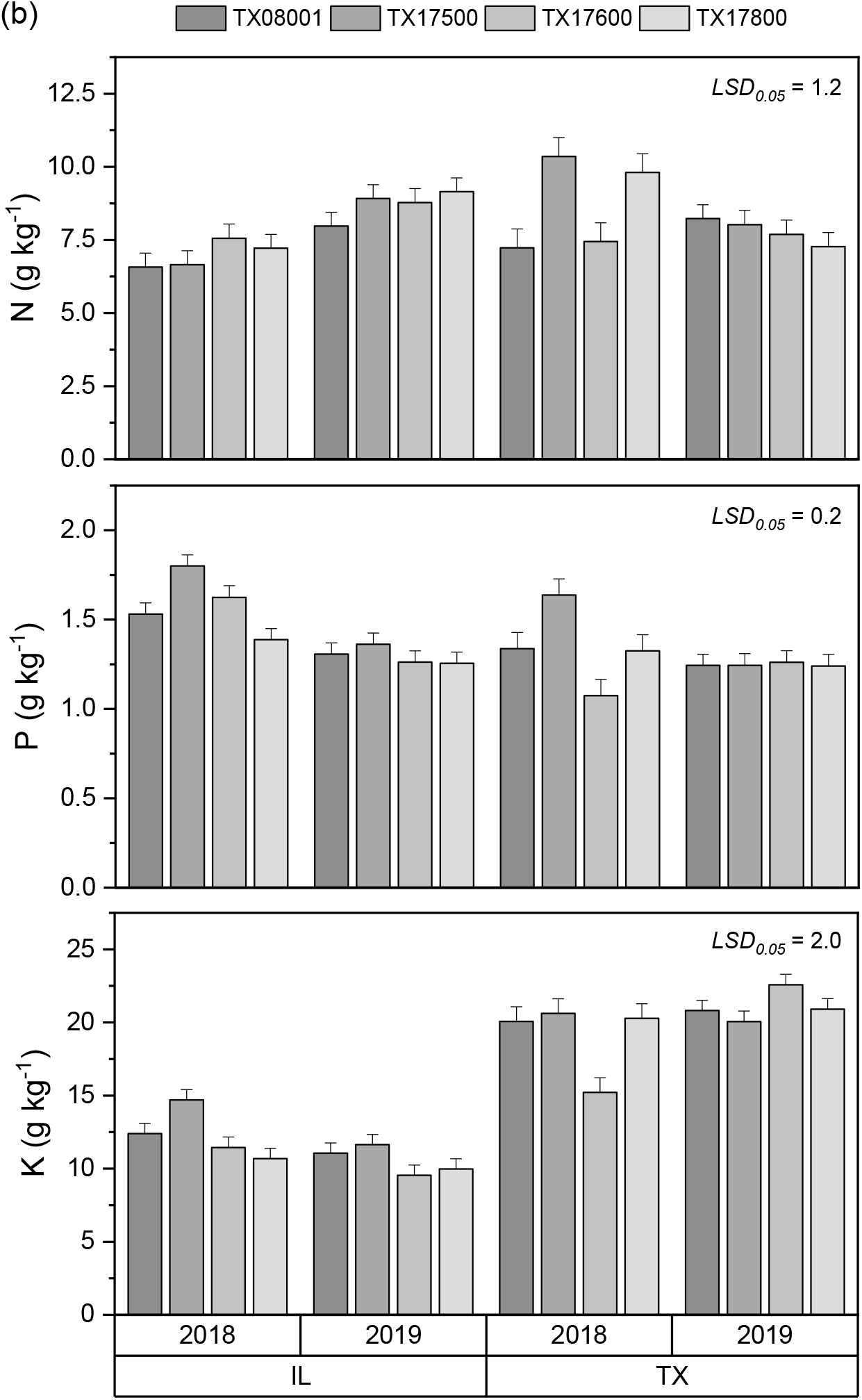
The significant effect of the 3-way interaction among location (IL and TX), year (2018 and 2019), and hybrid (TX08001, TX17500, TX17600, and TX17800) on (a) annual DM yields and (b) tissue nutrient (N, P, K) concentrations of energy sorghums.

### Nutrient Concentrations and Removal

The three-way interaction of location, year, and hybrids was also significant for tissue N, P, K concentrations (Table 4). In each environment, all hybrids showed similar tissue N concentrations except for biomass in TX-2018, which had higher variations in N concentrations (Fig. 3b). Averaged across the four hybrids N concentrations were generally higher in 2019 (8.7 g kg^−1^) than 2018 (7.0 g kg^−1^) in IL; by contrast, increased N concentrations were observed in 2018 (8.7 g kg^−1^) compared to the concentration in 2019 (7.8 g kg^−1^) in TX. This was likely due to the high N concentrations of the TX17500 (10.4 g kg^−1^) and TX17800 (9.8 g kg^−1^) hybrids in 2018 (Fig. 3b). For P concentrations, the TX17500 hybrid tended to have higher P concentrations than other hybrids in both locations, although differences were significant only in 2018. Across all environments, P concentrations in TX17500 biomass were approximately 13% higher than the average of the other three sorghum hybrids. All hybrids had higher tissue K concentrations in TX than in IL. The averages across all hybrids and the two years showed that the PS sorghum grown in TX increased the tissue K concentrations by approximately 75% (Table 5). For nutrient removal, the interaction among location, year, and hybrids only affected the biomass K removal, but the location x hybrids interaction influenced both P and K removal (Table 4). No consistent trend was observed for P and K removal among hybrids in both locations, but the average across hybrids and years showed that higher P removal occurred in IL (29.1 kg ha^−1^) than TX (23.9 kg ha^−1^). Conversely, the PS sorghum had a 63% higher K removal in TX compared to the sorghum grown in IL (Table 5).

Although the four hybrids had similar yield responses to the N rate (no interaction between N rate and other variables), the interaction of location and N rate was significant for the plant tissue N and K concentrations as well as their corresponding plant nutrient removal (Table 4). Both N and K concentrations responded differently to N rate in IL and TX. The tissue N concentration increased with increasing N rate in both locations (Table 6). Compared to the control zero N treatment, the highest N input of 168 kg ha^−1^ increased the N concentrations by approximately 38% in IL but only by 16% in TX (Table 6). By contrast, the 168 kg ha^−1^ N rate reduced the tissue K concentration by 14% compared to the 56 kg ha^−1^ N input, which was only observed in IL. The tissue P concentrations did not respond to N fertilizer rate. For nutrient removal, the ANOVA results also showed that the location x N rate interaction influenced the crop N and K removal (Table 4). The response of the biomass N removal to the increased N input was more substantial in IL than in TX. Compared to the zero N input, the 168 kg ha^−1^ N rate resulted in a 65% increase in N removal in IL but only 27% in TX (Table 6). Based on the linear relationship between N rate and removal, approximately an additional 0.5 kg ha^−1^ and 0.2 kg ha^−1^ of N were removed from the IL (R^2^ = 0.95) and TX (R^2^ = 0.81) soil, respectively, by increasing 1.0 kg-N ha^−1^ input. A significant response of K removal to N rate was only observed in TX. With an increasing N rate from 0 to 168 kg ha^−1^, the biomass K removal increased by nearly 22% with the removing rate of 0.5 kg K ha^−1^ per 1.0 kg-N ha^−1^ input (R^2^ = 0.90). No significant location x N rate effect on biomass P concentrations and removal was observed in this study.

### Nitrogen Use Efficiency

Three NUE indices (PNUE, NIE, and NRE) used in this study were also influenced by the environment. The location x year interaction was significant for PNUE and NIE. The highest PNUE appeared in the IL-2018 (151.3 kg kg^−1^), among others, but there was no year effect on PNUE in TX (Table 7). For the NIE, the IL location showed consistent NIE in both year (29.5 kg DM improvement per 1.0 kg-N input); however, approximately 19-fold differences between 2018 and 2019 was observed in TX (i.e., 2.6 kg DM in 2018 vs. 48.8 kg DM per 1.0 kg-N input shown in Table 7). The average across the two locations indicated that the NIE was higher in 2019 than 2018, so was NRE (18.1% in 2018 vs. 51.1% in 2019). The N effect on the three NUE indices, however, only influenced the PNUE. Although the location x N rate interaction showed a weak impact on PNUE at the P < 0.05 threshold (P = 0.0759), the PNUE reduced with increasing N input in both IL and TX (Table 7). Compared to the zero N treatment, the highest N rate (168 kg ha^−1^) resulted in declines in PNUE by around 25% and 17% in IL and TX, respectively. Neither the location x N-rate interaction or N-rate showed a significant impact on NIE and NRE, but the overall trend showed that the increased N input likely reduced both NIE and NRE.

**Table 7.**
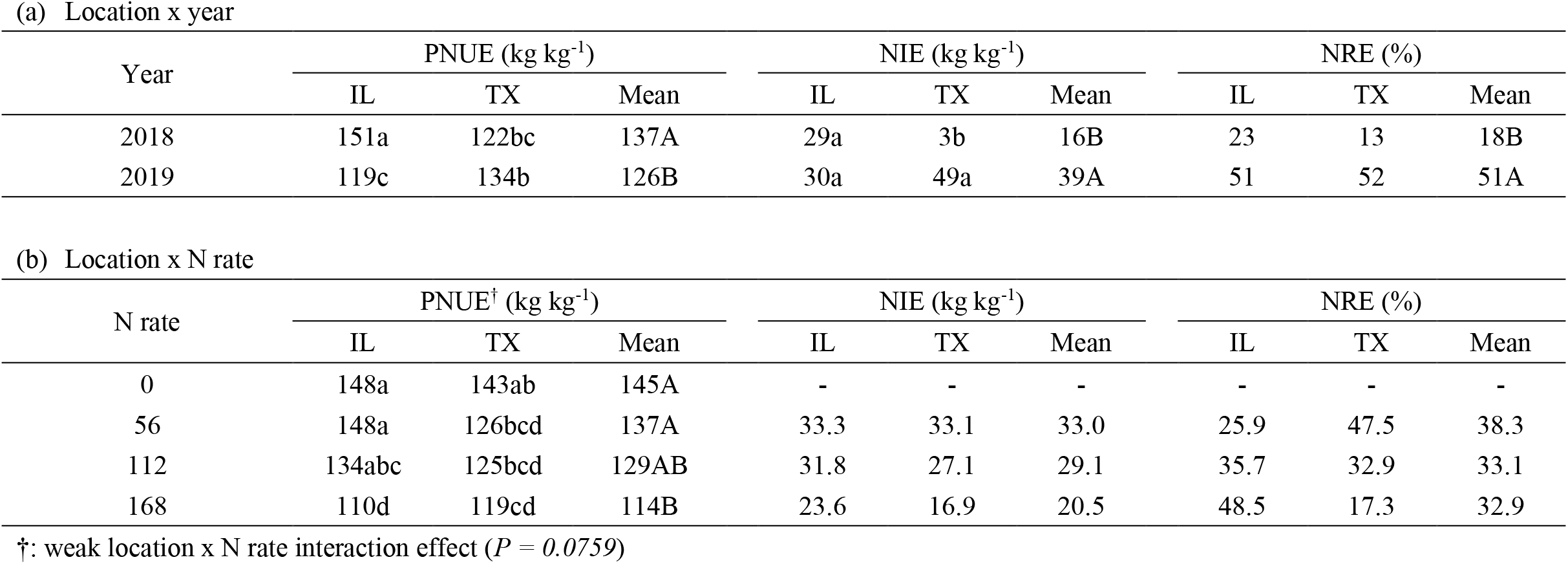
The significant 2-way interaction between (a) location (IL and TX) and year (2018 and 2019) and (b) location and N rate (0, 56, 112, and 168 kg-N ha^−1^) on indices of physiological N use efficiency (PNUE), N input efficiency (NIE), and N recovery efficiency (NRE). Lowercase and uppercase letters indicate mean separation α=0.05 organized highest to lowest value for year x location and year, respectively (no mean separations were applied if the variable effect was not significant).

### Chemical Composition

Sorghum feedstock chemical composition was influenced by the environment, hybrid, N rate, and their interactions (Table 8). The 4-way interaction of location, year, N-rate, and hybrid was not significant for the biomass structural and soluble compositions; however, the 3-way interaction among location, year, and hybrid showed a significant impact on the concentrations of structural and soluble components (Table 8). All hybrids in TX-2019 had the lowest structural component (a sum of glucan, xylan, lignin, galactan, arabinan, acetyl, and protein) concentrations among three environments (IL-2018, IL-2019, and TX-2018) and TX17600 tended to have the lowest structural component concentration among all hybrids (Fig. 4). Two VPS hybrids (TX08001 and TX17500) showed similar compositions within a growing environment, but the concentrations differed across environments (Fig. 4). Across two experimental years, the effect of the 2-way interaction between locations and hybrids on the biomass structural and soluble components are shown in Table 9. For both IL and TX, the TX17800 hybrid consistently contained higher concentrations of the structural glucan, xylan, and lignin than other hybrids; likewise, TX17800 tended to have higher concentrations for both biochemical- and thermochemical-processing interested components (i.e., BIC and TIC). For instance, the BIC and TIC concentrations of the TX17800 hybrid was approximately 5 % and 7 % higher, respectively, compared to other hybrids (Table 9). In contrast, TX17800 had the lowest concentrations of soluble sucrose and other non-structural inorganics (e.g., ash). TX17600 had the highest sucrose concentrations among hybrids by increases of approximately 34 % and 89 % in IL and TX, respectively (Table 9). For other structural components, the TX08001 (VPS hybrid) had the highest concentrations of galactan, and the TX17800 (MPS hybrid) had the lowest arabinan among hybrids. The VPS hybrid TX17500 had the highest biomass ash concentration among hybrids. Between two locations, the TX site generally had higher concentrations of the BIC, TIC, acetyl, corresponding to the lower ash concentrations than observed at the IL site (Table 9). The N effect on feedstock composition showed that only a few chemical compositions, including lignin, acetyl, ash, and sucrose, responded significantly to the N rate for all hybrids and no interaction of N rate x hybrid was observed (Table 8). The increased N rate increased lignin and ash concentrations but reduced acetyl and sucrose concentrations (Table 10).

**Table 8.**
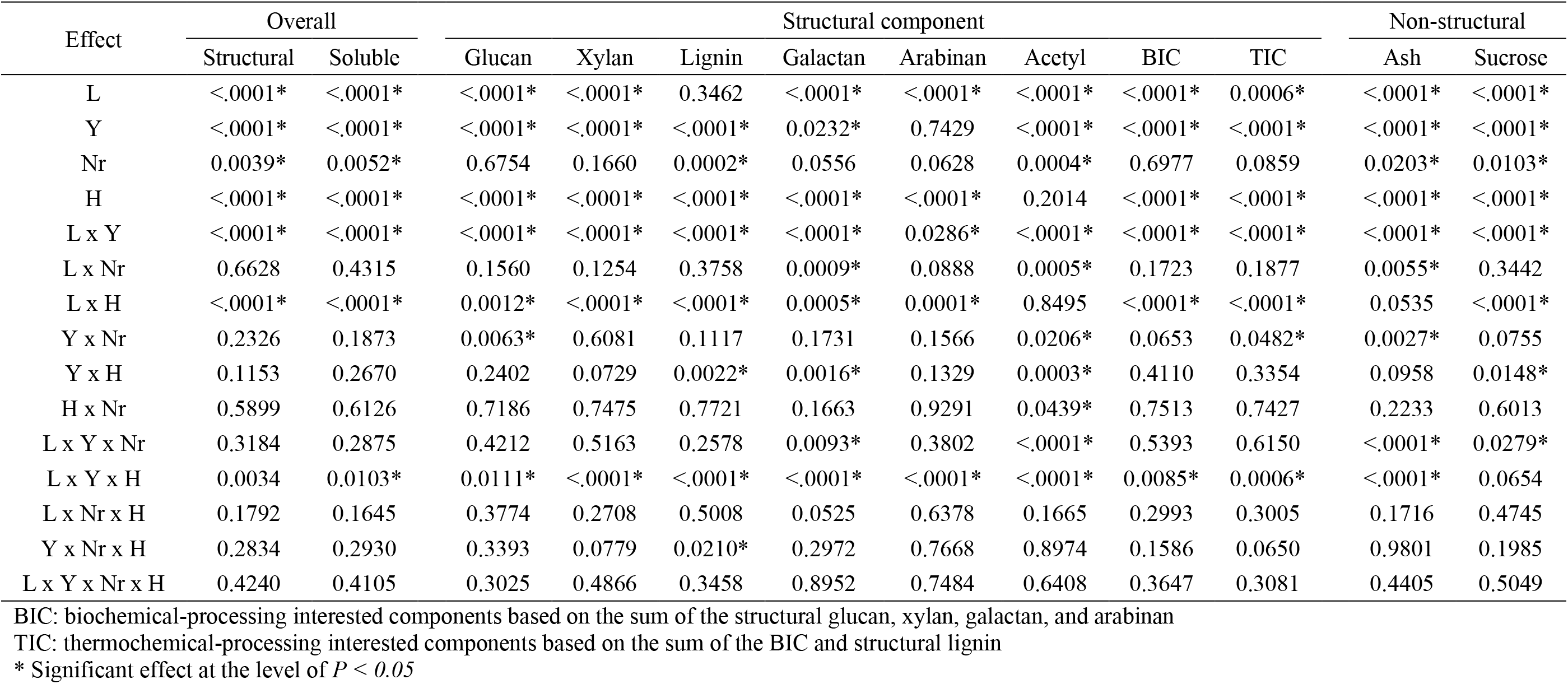
Analysis of variance (ANOVA) showed the effects of main factors of location (L), year (Y), N-rate (Nr), and hybrids (H) and their interactions on feedstock chemical compositions with a significance level of 0.05.

**Table 9.** The significant effects of location (IL and TX) and hybrid (TX08001, TX17500, TX17600, and TX17800) on concentrations of feedstock components. Lowercase letters indicate mean separation α=0.05 organized highest to lowest value (no mean separations were applied if the variable effect was not significant).

| Location | Hybrid | Structural components (g kg<sup>-1</sup>) |  |  |  |  |  |  |  | Non-structural (g kg<sup>-1</sup>) |  |
| --- | --- | --- | --- | --- | --- | --- | --- | --- | --- | --- | --- |
|  |  | Glucan | Xylan | Lignin | Gal | Arab | Acetyl | BIC | TIC | Ash<sup>†</sup> | Sucrose |
| IL | TX08001 | 285.4ef | 166.2a | 115.7c | 11.5b | 25.9b | 12.5 | 489.0d | 604.7d | 94.3b | 57.7de |
|  | TX17500 | 279.3f | 162.7b | 108.0d | 11.1c | 26.0b | 12.1 | 479.1e | 587.1e | 99.6a | 62.3cd |
|  | TX17600 | 292.0de | 153.8d | 115.6c | 10.7d | 24.2c | 12.6 | 480.5e | 596.2de | 90.6b | 73.1b |
|  | TX17800 | 305.5b | 168.2a | 126.8a | 10.7d | 22.7d | 12.3 | 507.1b | 633.9b | 93.7b | 43.8f |
| TX | TX08001 | 297.2cd | 159.0c | 120.0b | 11.9a | 27.7a | 16.8 | 495.9cd | 615.9c | 85.6d | 65.9c |
|  | TX17500 | 299.1bc | 163.1b | 118.7bc | 11.5b | 27.0a | 16.5 | 500.7bc | 619.4c | 86.8c | 56.6de |
|  | TX17600 | 292.3d | 142.4e | 102.4e | 11.6b | 27.6a | 16.9 | 473.9e | 576.2f | 81.3e | 110.3a |
|  | TX17800 | 315.5a | 166.1a | 128.4a | 11.5b | 26.0b | 17 | 519.3a | 647.5a | 82.9d | 51.9e |
| Hybrid mean | TX08001 | 291.3b | 162.6b | 117.9b | 11.7a | 26.8a | 14.6 | 492.4b | 610.3b | 90.0b | 61.8b |
|  | TX17500 | 289.2b | 162.9b | 113.3c | 11.3b | 26.5ab | 14.3 | 489.9b | 603.2b | 93.2a | 59.4b |
|  | TX17600 | 292.1b | 148.1c | 109.0d | 11.2bc | 25.9b | 14.8 | 477.2c | 586.2c | 86.0c | 91.7a |
|  | TX17800 | 310.5a | 167.2a | 127.6a | 11.1c | 24.4c | 14.6 | 513.2a | 640.7a | 88.3b | 47.9c |
| Location mean | IL | 290.5b | 162.7a | 116.5a | 11.0b | 24.7b | 12.4b | 488.9b | 605.4b | 96.0a | 59.2b |
|  | TX | 301.0a | 157.6b | 117.3a | 11.6a | 27.1a | 16.8a | 497.5a | 614.8a | 82.6b | 71.2a |

Gal: Galactan
Arab: Arabinan
BIC: biochemical-processing interested components based on the sum of the structural glucan, xylan, galactan, and arabinan
TIC: thermochemical-processing interested components based on the sum of the BIC and structural lignin
†: weak location x hybrid interaction effect ( $P = 0.0535$ )

**Table 10.**
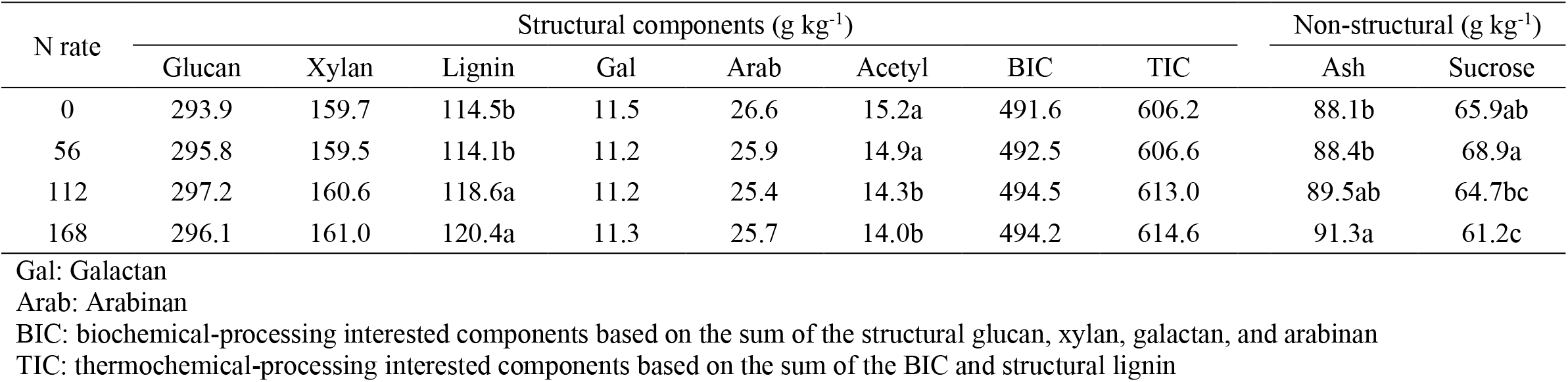
The significant effects of N rate (0, 56, 112, and 168 kg-N ha^−1^) on concentrations of feedstock composition. Lowercase letters indicate mean separation α=0.05 organized highest to lowest value (no mean separations were applied if the variable effect was not significant).

**Figure 4.**
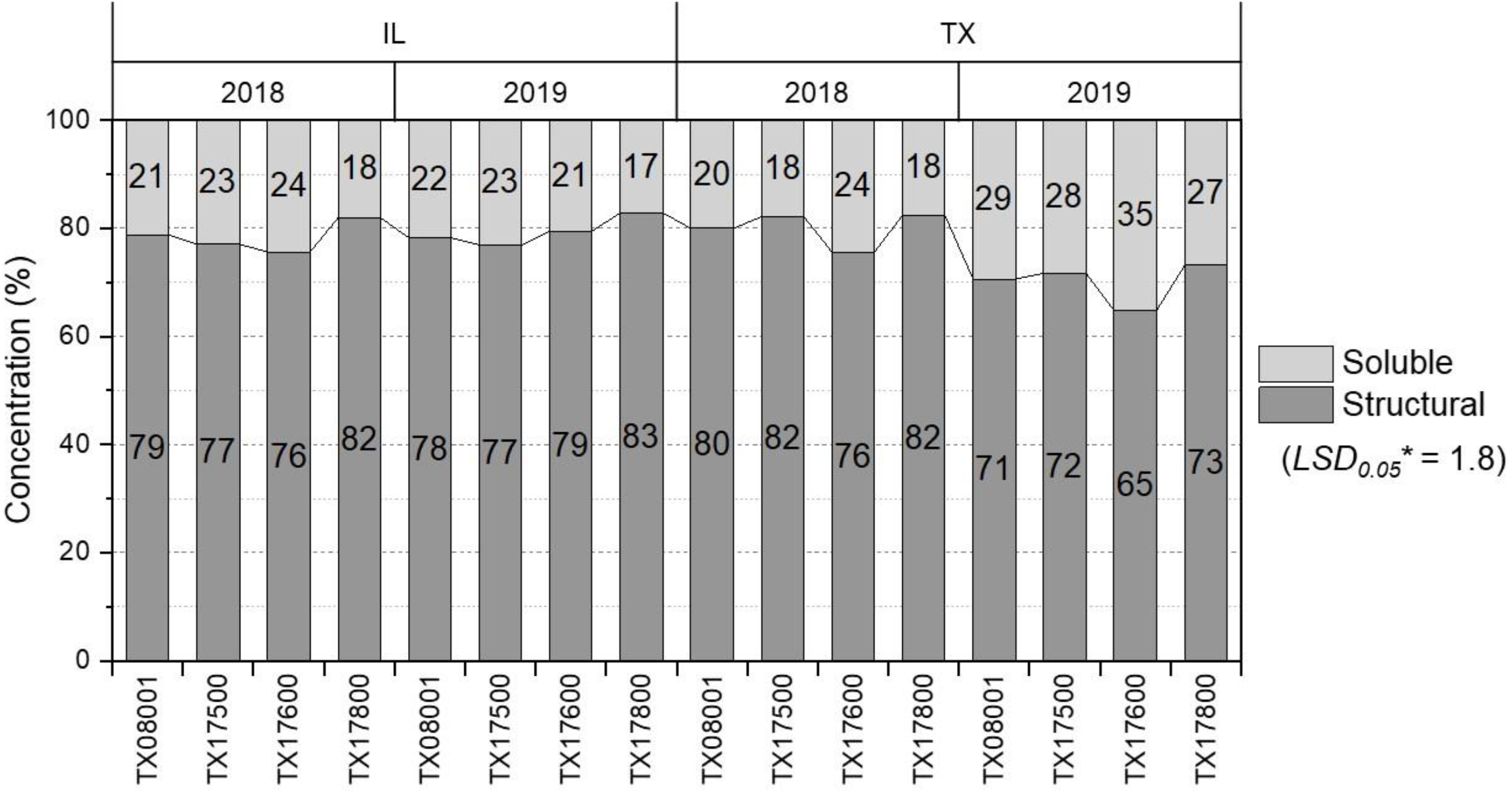
The significant 3-way interaction between location (IL and TX), year (2018 and 2019), and hybrid (TX08001, TX17500, TX17600, and TX17800) and their influence on concentrations of dry biomass structural and soluble compositions of energy sorghums (*Both structural and soluble concentrations had the same Fisher’s least significant difference [LSD] at α=0.05 because the measured concentrations were normalized to the percentage bases).

## Discussion

### Environment Effects

The cultivation environment plays a significant role in biomass yield, nutrient concentrations and removal, feedstock chemical compositions, and the strategies for optimizing management practices [3,11,20,21,23,37]. Four environmental regimes (two locations by two years) were evaluated for the yield potential, biomass nutrient concentrations and removal, and feedstock compositions of four PS sorghum hybrids in this study. The PS nature of these hybrids keeps them in the vegetative stage and they never reach the reproductive stage. This allows them to continuously accumulate biomass yield by increasing both leaf area index and solar radiation inception during the growing season [5,7,11,38]. The 2-year-average yield of two VPS hybrids (TX08001 and TX17500) grown in the IL site increased by 10% and 22%, respectively, compared to the TX site (Table 5). Increased biomass yield was attributed to several potential factors including longer days at the peak of the growing season, cooler night temperatures and more consistent rainfall in the growing season (Table 1). Two MPS hybrids (TX17600 and TX17800) responded differently to the growth period/cumulative daylength. The TX17600 hybrid exhibited higher yield potential (2-year average) in TX (22.5 Mg DM ha^−1^) than IL (19.5 Mg DM ha^−1^), which was possibly due to the prolonged growth period in TX (Table 1); by contrast, the TX17800 hybrid showed higher yield potential in IL.

Weather variations also influenced biomass yield significantly [10]. The monthly temperature in IL and TX were consistent in both the experiment years compared to the 30-year averages; however, the precipitation was highly variable (Fig 2). The great variability in precipitation not only raises the difficulties for optimizing the field management but also influences crop development as well as biomass productions [39–41]. In TX, for instance, the seasonal precipitation was concentrated during the late growing season (Sept. and Oct.) in 2018; whereas, most of the seasonal precipitation accumulated during the early season (May to July) in 2019 (Fig. 2). The biomass yields across the two years showed more variation in TX than IL, likely resulting from the highly fluctuating precipitation pattern (Fig. 3). For instance, both VPS hybrids and TX17800 had lower biomass yields in the TX-2018 regimes than in the other three environments. This yield decline was attributed to the drought condition during the early growing season from May to August (2018: 150-mm vs. 30-year average: 324-mm) in TX, 2018. Inadequate rainfall slows the crop’s early establishment and delays growth and yield [21,40]. In this study, the overall biomass yield of the PS sorghum was similar to our previous results reported by Maughan et al. (2012) [10], but approximately 30% lower than the other reports (>30 Mg DM ha^−1^) where the field trials were mostly in the long-growing season (> 150 days) areas [11–13]. The relatively low yield was presumably due to the short production seasons and can be improved by prolonging the growing season, such as early planting [7].

A substantial environmental impact on tissue nutrient concentrations shown in this study was also observed in several studies [29,34,42–45]. The decline in N concentrations was usually corresponding to the increase in biomass yield, which was attributed to the effect of a growth dilution [42,45,46]. This dilution effect on P and K concentrations, however, was not observed in this study, which was also reported by Maw et al. (2020) [45]. Compared to IL, the higher soil K content in TX, showing approximately 4-fold soil K (Table 2), likely resulted in an increase in biomass K concentrations for all hybrids (Table 5). The increased concentrations of the alkali metals, such as K and Na, in biomass substantially affect energy conversion efficiency from the feedstock [15,45,47]. The PS sorghum grown in TX also led to an approximately 34% increase in biomass Na concentrations compared to the sorghum from IL (data not shown). The increased K concentration also resulted in higher K removal in TX than IL. Since nutrient removal is the product of biomass yield and tissue nutrient concentrations, the reduced yield (average across four hybrids) led to lower K removal in 2018 than 2019 in TX and the lowest P removal among for environmental regions (IL-2018: 32.9 kg ha^−1^; IL-2019: 25.3 kg ha^−1^; TX-2018: 21.0 kg ha^−1^; TX-2019: 26.7 kg ha^−1^; *LSD*_0.05_ = 2.4).

Feedstock compositions were also influenced by the grown environment [12,16,20]. As lignocellulosic bioenergy materials, it is crucial to minimize the content of the non-structural components but water stress has been reported to increase the biomass non-structural components for different bioenergy feedstocks, such as corn stover, miscanthus, switchgrass, and mixed perennial grasses [48–50]. McKinley et al. (2018) [12] reported that the soluble compounds in energy sorghum stem biomass were generally higher, along with increased sucrose, in rainfed systems than irrigated systems. For these four hybrids, a significantly increased concentration of the soluble components was observed in the TX-2019 regimes (Fig. 4).Dry conditions in the TX-2019 began in July as is typical compared to the 30-year benchmark (Fig. 2). Since the TX-2019 site received adequate rain early to facilitate crop establishment, the following dry season (Jul. to Oct.) did not affect the biomass yield of the well-established sorghum due to a good water stress resistance; however, the non-structural compounds did respond to the dry environment [50].

### Hybrid Comparisons

Among four sorghum hybrids, no consistent yield trends were observed in IL. In TX, the TX17600 hybrid consistently produced higher biomass yield than the other hybrids (Fig.3). The TX17600 hybrid also showed less sensitivity to the environmental variation for biomass productions, resulting in the mean DM yield of 21±2 Mg ha^−1^ across the four cultivation regimes. For the nutrient concentration, the four PS hybrids showed similar aboveground biomass N concentrations under similar cultivation environments besides the adverse condition in the TX-2018 region (limited precipitation in the early growing season) for crop establishment (Fig. 3b). Excluding the data from the TX-2018 environment, the ranges of the biomass N concentrations (6.6 and 9.2 g kg^−1^) and the corresponding yields (19.2 to 23.6 Mg DM ha^−1^) were similar to the previous studies on the fiber, sweet, and biomass sorghums under a similar N input [34,42,44]. Other PS sorghum studies, conversely, reported lower N concentrations (4.4 ~ g kg^−1^) associated with the higher biomass yield (> 25 Mg DM ha^−1^) [31,43]. The improved yield, usually resulting from an extended growth period (e.g., > 140 growth days), diluted the biomass N concentrations [31,42,43]. Although the four PS hybrids had similar N requirements, the TX17500 showed a higher stover P concentration/requirement than the other hybrids. The high biomass sorghum hybrids with lower tissue nutrient concentrations, usually along with lower nutrient removal, have more sustainable benefits for the cropping system [45,51].

The composition analysis (Fig. 4 and Table 9) indicated that the TX17600 hybrid had a juicier stalk and higher sucrose concentration than other hybrids, and its environmentally-insensitive and MPS characteristics are likely favorable for stable and high yield productions by growing TX17600 in the region with a more extended harvest season (e.g., subtropical or tropical areas) [3]. Understanding the feedstock compositions are essential for both qualities and quantities of the biofuel products, and different quality attributes can be used as indices based on the conversion technology [15,52]. Among hybrids, the TX17800 hybrid consistently resulted in the highest concentrations of the energy-rich components mainly contributed by the glucan, xylan, and lignin. In temperate region such as IL, the TX17800 hybrids can be considered a good candidate for lignocellulosic energy crops using either bio- or thermo-processes because of the excellent yield potential along with high feedstock quality [15]. In tropical or subtropical regions of TX, the TX17600 produced steady yield with high carbohydrate-rich compounds (Table 9). The two VPS hybrids (TX08001 and TX17500) in IL did not show distinct differences in terms of yield potential (Table 5); however, the TX08001 hybrid showed a higher feedstock quality than TX17500 by increasing the energy potential (higher BIC and TIC concentrations) and lowering ash concentrations (Table 9). The first-generation TX08001 energy sorghum hybrids have been consistently reported for both high yield and quality potentials in several studies [7,12,31]. The acetic acid, derived from the acetyl functional group, can be produced during the hydrolysis processes (e.g., the dilute acid or enzymatic hydrolysis pretreatment), and the increased acetic acid likely inhibit the fermentation effectiveness (biological inhibitor) and corrode the processing pipelines [52,53]. In both locations, no differences in acetyl concentration were observed among hybrids; however, the sorghum grown in TX resulted in higher acetyl concentrations than IL (Table 9).

### Nitrogen Effects and Use Efficiency

Nitrogen, a primary nutrient for synthesizing amino acids, nucleic acids, or other essential organic compounds, facilitates crop growth. Although the four hybrids and two locations showed similar yield responses to different N inputs in this study, the increased N supply generally improved biomass yield and the standout N rate for significant yield improvement was less than 120 kg-N ha^−1^, which was also shown in other studies [4,20,26,54,56,57]. This study showed that increased N input led to an increase in biomass N concentrations, which was often reported in many studies [34,44,56,58]. The improved biomass yield due to N supply can dilute P and K in plant tissue and lower their concentrations, which was only observed for tissue K concentrations in IL [44,45,56]. The limited response of P and K concentrations to N input was presumably due to 1) the substantial environmental impact on tissue nutrient concentrations, and 2) a sufficiency supply of P and K nutrients in the soil for plant uptake [45,59]. For nutrient removal, plant N removal increased with more N input because of the increases in both biomass yield and N concentration [34,44,56]. Compared to TX, the increased N removal rate in IL (~0.5 kg N ha^−1^ uptake per 1.0 kg N input) was due to higher yield and tissue concentration responses to the N fertilizer. The N removal in TX (~0.2 kg N ha^−1^ uptake per 1.0 kg N input) was similar in several reports [29,34,56]. In TX, the increased K removal with increasing N input was a result of the DM biomass accumulation [45]. The feedstock compositions were less influenced by different N rates compared to the biomass yield, which is similar to several studies [20,57,60,61]. The reduced sucrose concentrations occurring at a high N rate in this study (Table 10) were also observed by Almodares et al. (2009) [26]. With increasing N input, the improved lignin content potentially increased the yields of biofuel products, such as crude oil; however, the increased ash content might lead to an adverse impact in the conversion effectiveness [15,62].

Nitrogen use efficiency can be evaluated based on the biomass DM accumulation per unit of N in the plant tissue (PNUE), and responses of biomass yield and crop nutrient removal to the applied N fertilizer (NIE and NRE) for optimizing N management [31,34,56,63]. Variations in the growing environment can be the main factor to influence NUE as well as the associated management strategies for the energy sorghum cultivation [34,56]. In IL, the increase in PNUE in 2018 was due to the improved DM yield corresponding to the lowered biomass N concentrations, which was likely due to a favorably growing condition, such as a sufficient water supply [34]. Likewise, the severe water stress during the early growing season delayed crop development and limited the yield response to N supply, resulting in the lowest NIE in the TX-2018 region. For both locations, favorable weather conditions also led to higher NRE in 2019 than in 2018, resulting in 51% NRE in 2019. This study showed a similar range of PNUE (78.1 – 256.4 kg DM kg^−1^) to the reports from Grennel et al. (2014) [64] and Maw et al. (2017) [34], and the PNUE likely improved with the low N input [34,65,66]. Although the N effect on NIE and NRE was not significant in this study, the better NIE and NRE likely occurred at the low N-rate [34,56,65,66,69]. Many studies have shown that the biomass sorghum has a low N fertilizer requirement (mostly < 120 kg-N ha^−1^) for optimizing biomass yield with desirable feedstock quality for biofuel productions [29,56,57,67–69]. Moreover, the biomass yield, PNUE, and NIE can be further improved by extending the growing season [11,12,31]. Even though the low N input was favorable in this study, the adequate N supply could facilitate PS energy sorghum to achieve its maximum yield potential and nutrient use efficiency by providing the best growing conditions (e.g., irrigation, early planting for extending season length) for the optimum biomass production and N accumulation [4,7,12,35].

## Conclusion

This study found cultivation location, year, and hybrid significantly influences biomass yield of photoperiod-sensitive sorghum. The 2-year-averaged yield of two VPS sorghum hybrids (TX08001 and TX17500) grown in the IL site was higher by 10% and 22%, respectively than the TX site. Compared to TX, the more consistent weather pattern in IL resulted in more consistent performance in biomass yield among hybrids, which is favorable for a stable feedstock production and nitrogen (N) management. The TX17600 performed consistently in biomass productions with the mean annual yield of 21±2 Mg DM ha^−1^ and feedstock compositions over the four growing environments displaying resilience to adverse conditions. The four sorghum hybrids showed similar yield responses to N rate, and the lower N input (> 112 kg-N ha^−1^) tended to have a higher N use efficiency in both locations. The TX17800 hybrid consistently resulted in the highest concentrations of the energy-rich components mainly contributed by the glucan, xylan, and lignin. It is important to consider the end-use conversion process, i.e., thermo-chemical, biochemical, when selecting hybrids. The location and growing season environment also influenced chemical compositions. Drought stress increased non-structural components in the biomass. This increase in non-structural components possibly decreases conversion efficiencies for numerous biomass conversion techniques. Testing hybrids in the regions of conversion facilities would be beneficial to the biofuels industry as results varied with the other hybrids in the trial. Bioenergy sorghum is a viable annual biomass feedstock for the Midwest corn-belt; as well as the cotton-belt. Drought tolerance, low N requirement, and biomass yields of 20 Mg DM ha^−1^ are all benefits of the adoption of bioenergy sorghum.

## Acknowledgements

Author #1 (AS) and Author #2 (CHL) have equally contributed to this study. This work was funded by the DOE Center for Advanced Bioenergy and Bioproducts Innovation (U.S. Department of Energy, Office of Science, Office of Biological and Environmental Research under Award Number DE-SC0018420). Any opinions, findings, and conclusions or recommendations expressed in this publication are those of the author(s) and do not necessarily reflect the views of the U.S. Department of Energy.

